## Supplementary material for "Connecting metapopulation heterogeneity to aggregated lifetime statistics"

Eduardo H. Colombo[1, *]

[1] IFISC (CSIC-UIB), Campus Universitat Illes Balears, 07122, Palma de Mallorca, Spain

### I. APPENDIX S1: RELATION BETWEEN LIFETIMES AND PATCH PROPERTIES

In this section, we provide two examples of mathematical models for the population dynamics in order to establish a link between the local average lifetime $\tau = \langle T_i \rangle$ and the patch property $m$.

In the following developments, we introduce the standard ingredients, such as logistic growth rate and demographic and environmental fluctuations, that constitute the canonical model [1, 2]. Following the population density approach, we write the set of coupled ordinary differential equations for the patches' population densities $u_i$,

$$\dot{u}_i = f_i(u_i) + \sqrt{u_i}\xi_i(t) + u_i\eta_i(t) + D\sum_{j\neq i}\gamma(|x_i - x_j|)u_j \,. \tag{1.1}$$

for $i = 1, 2, \ldots, N$.

The local dynamics is given by $f_i$ plus the demographic and environmental fluctuations, which are commonly approximated by zero mean white Gaussian noises with variances $\sigma_\xi^2$ and $\sigma_\eta^2$, respectively. Nevertheless, environmental fluctuations receive an special treatment, considering that small correlation are present, being handled within the Stratonovich prescription [1, 3, 4]. For $f_i(u_i) = r_i u_i(1 - u_i/K_i)$, being $r_i$ the local growth rate and $K_i$ the carrying capacity, it has been considered that local part of Eq. (1.1) constitute a canonical model for population dynamics [1–3]. The last term of Eq. (1.1) couples the subpopulations, representing the fluxes between patches with rate $D$ and intensity that depends on the distance through $\gamma$.

#### A. Effective model

The effective model arises when we bypass the challenges of solving the full set of stochastic differential equation [5], setting by hand the incoming flux of individuals at each patch. When the metapopulation has passed any transient dynamics and a quasi-stationary state has been achieved, in the perspective of each subpopulation $i$ we can attribute an incoming flux of individuals $\phi_i(t)$, such that we approximate

$$D\sum_{j\neq i}\gamma(|x_i - x_j|)u_j \sim \phi_i(t) \,. \tag{1.2}$$

*

The effective model for each patch is then defined by

$$\dot{u}_i = f_i(u_i) + \sqrt{u_i}\xi(t) + u_i\eta_i(t) + \phi_i(t) \,. \tag{1.3}$$

Given the function $f_i$ and the migration protocol $\phi_i$, it is possible to obtain the patch lifetime $\tau_i$ distribution which is a function of the local properties, e.g the carrying capacity or migration rates [1–3].

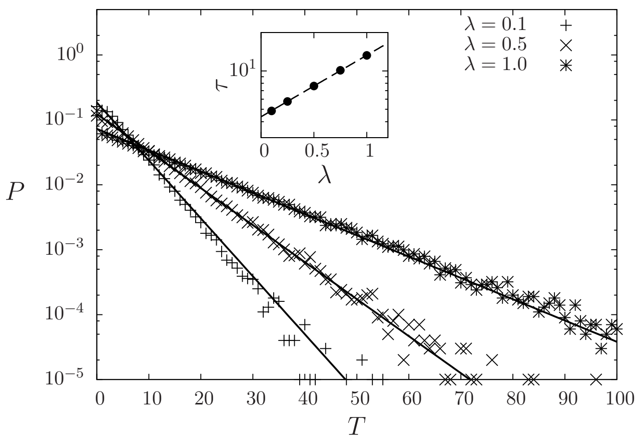


Figure 1. Probability distribution of patch lifetimes $P$ for different average migration rates $\lambda$ for Eq. (1.3) with $r_i = K_i = 1$. The solid lines represent the approximation $\exp(-T/\tau)/\tau$, considering the migration term $\phi$ as a Poisson shot noise with one unit fixed pulse amplitude. In the inset, we show the behavior of $\tau(\lambda)$, where the dashed line represent an exponential fit.

In Fig. 1, we present the lifetime probability distribution for a given patch with different migration rates $\lambda$ assuming a Poisson shot noise protocol

$$\phi(t) = \sum_i \delta(t - t_i) \,, \tag{1.4}$$

where the migration inter-event times $t_{i+1} - t_i$ are exponentially distributed with mean $1/\lambda$. The exponential law exhibited by Eq. (1.3) is robust against different modeling considerations [6] and a key feature for the developments in the deconvolution operation. In the inset of Fig. 1, we show that the mean lifetime increases exponentially with the migration rate $\tau(\lambda) \sim e^\lambda$, also a typical result found in the literature, not only for the migration rate, but also for the carrying capacity [2]. Analytically, these results can be extracted from approximative expressions based on the theory of first-passage times [1–3, 7].

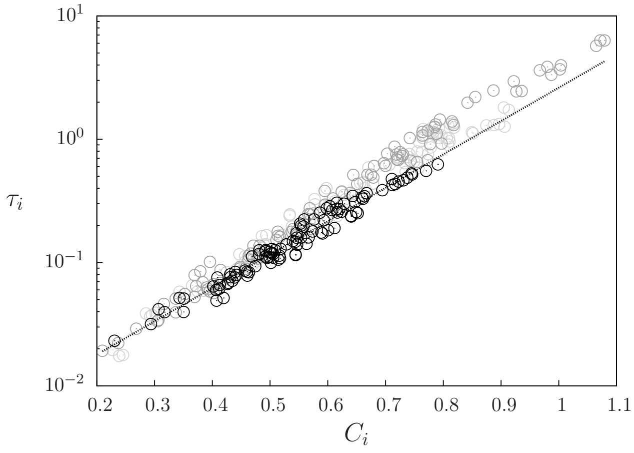


Figure 2. Mean lifetime $\tau$ vs patch connectivity $C$ for a random landscape in a $10 \times 10$ domain with periodic boundary conditions, in which 100 patches are distributed randomly with $r_i = K_i = 1$. The black dots represent cases with $D = 0.01$ and the gray dots $D = 1$.

For the cases in which $\phi$ is described by a Gaussian noise, standard first-passage times techniques find expressions for the mean lifetime which are able to reproduce the form $\tau \sim e^{\langle\phi\rangle} \sim e^K$ used in the application section [2, 8]. However, it has been shown that the use of more sophisticated techniques can improve the precision in cases in which the Fokker-plank approach fails, for instance, when assuming strong and correlated environment fluctuations [2, 9, 10]. In extreme cases, the mean lifetime might follow a power-law [2].

#### B. Spatial explicit model

A similar result from the one in I A can be extracted directly from the spatial explicit model. Analogously, here, we find that the mean patch lifetime grows exponentially with connectivity, that being the counter part of migration rate in the effective model. In this section we solve numerically the spatial explicit model given by Eq. (1.1) assuming a random landscape (randomly distributed patches under periodic boundary conditions) and $\gamma(r) = e^{-r}$ [8].

Defining that the patch connectivity $C_i$ is given by

$$C_i \equiv \sum_{i \neq j} \gamma(|x_i - x_j|) \,, \tag{1.5}$$

in Fig. 2, we show the simulation results for each patch mean extinction times as a function of its connectivity $C_i$. For this case, the lifetime probability distribution also follows an exponential law with mean extinction time that grows with $C$. It is relevant to note that, even for strongly coupled cases, the exponential law is preserved.


[1] H. Hakoyama and Y. Iwasa, Journal of Theoretical Biology **232**, 203 (2005).
[2] O. Ovaskainen and B. Meerson, *Trends in Ecology & Evolution*, Trends in Ecology & Evolution **25**, 643.
[3] H. HAKOYAMA and Y. IWASA, Journal of Theoretical Biology **204**, 337 (2000).
[4] L. A. da Silva, E. H. Colombo, and C. Anteneodo, Phys. Rev. E **90**, 012813 (2014).
[5] O. Ovaskainen, D. Finkelshtein, O. Kutoviy, S. Cornell, B. Bolker, and Y. Kondratiev, Theoretical Ecology **7**, 101 (2014).
[6] V. Grimm and C. Wissel, Oikos **105**, 501.
[7] N. Van Kampen, *Stochastic Processes in Physics and Chemistry*, North-Holland Personal Library (Elsevier Science, 2011).
[8] E. H. Colombo and C. Anteneodo, Phys. Rev. E **92**, 022714 (2015).
[9] A. H. O. Wada, M. Small, and T. Vojta, Phys. Rev. E **98**, 022112 (2018).
[10] A. Kamenev, B. Meerson, and B. Shklovskii, Phys. Rev. Lett. **101**, 268103 (2008).
